# Human dental pulp stem cells grafted into C57BL/6J hippocampus differentiate towards immature neuronal like cells displaying action potential firing activity

**DOI:** 10.64898/2026.06.16.732586

**Authors:** Beatriz Pardo-Rodríguez, Irene Manero-Roig, Jone Salvador-Moya, Ruth Basanta-Torres, Daniel Martín-Aragón, Sandra Hernández-Sánchez, Jon Luzuriaga, Ander Martín, Aurélie Lampin-Saint-Amaux, Frédéric Lanore, Fernando Unda, Gaskon Ibarretxe, Jose Ramon Pineda

**Author notes:** These authors contributed equally to this work (Gaskon Ibarretxe, Jose Ramon Pineda).

## Abstract

Stem cell therapy represents a promising strategy for the replacement and functional restoration of damaged neural tissue in neurodegenerative conditions. Human dental pulp stem cells (hDPSCs) have emerged as potential candidates for neuroregeneration due to their ease of isolation, neural crest origin, neurotrophic and anti-inflammatory capacity, and demonstrated ability to differentiate *in vitro* into neuronal-like cells exhibiting electrophysiological activity. Although the immunomodulatory and neuroprotective properties of hDPSCs have been reported in multiple models of brain disease, their capacity to functionally integrate into host neuronal circuits remain poorly understood. In this study, we have grafted green fluorescent protein (GFP)-transduced, neural preconditioned hDPSCs into the CA1 region of the hippocampus of C57BL/6J mice. One month after transplantation, GFP^+^-hDPSCs survived in the brains of non-immunosuppressed mice and remained localized within the grafted area. Notably, the transplanted cells underwent *in situ* differentiation and exhibited a neuroblast-like phenotype, characterized by positive doublecortin expression and immature neuronal-like electrophysiological properties, like high membrane input resistance, low capacitance, and the ability to generate single action potentials after stimulation. Together, these findings provide the first evidence that hDPSCs can survive and integrate into the hippocampal network of the mouse brain at one-month post graft, supporting their potential use for future therapeutic applications in acute brain lesions and neurodegenerative disorders.

---

Hippocampus, Cell therapy

## Background

The rapid ageing of the global population is accompanied by a marked increase in the incidence of neurodegenerative diseases (1,2). Current therapeutic strategies for neurological diseases largely focus on symptomatic relief and slowing disease progression, rather than addressing the underlying pathology. Likewise, rehabilitation approaches can enhance functional recovery but do not directly replace lost neural tissue (3,4). Even though numerous neuroprotective drugs and gene-based therapies have been developed, their clinical translation remains challenging and their long-term efficacy is often limited (5,6). These limitations have prompted a growing interest in alternative approaches, including stem cell-based therapies, which can target disease pathophysiology through multiple mechanisms such as immunomodulation, cell replacement, paracrine signaling, and activation of endogenous repair processes (7).

Within this framework, human dental pulp stem cells (hDPSCs) have emerged as a particularly promising source due to their ease of extraction, lack of ethical concerns, and neural-related embryological origin (8). Being neural crest derived cells, the differentiation capacity of hDPSCs extends beyond mesenchymal cell lineages to include neural progenitors and neuronal-like cells expressing mature neuronal markers, without a need for genetic manipulation (9–12). In addition to their differentiation capacity, hDPSCs exhibit strong paracrine activity in the context of brain injury, secreting immunomodulatory cytokines (13) and neurotrophic factors (11,14). These properties may contribute to neuroprotection of the injured tissue and attenuation of the chronic inflammatory responses that drive progressive neural loss following brain damage.

To define a cell as a fully differentiated functional neuron, it must not only express neuron specific gene products but also exhibit neuron specific electrophysiological activity, including tetrodotoxin (TTX)-sensitive action potentials (APs) (15). Until now, evidence supporting the electrophysiological properties of neural-differentiated hDPSCs remained limited. Only two studies, published by Gervois et al. in 2015, and Li et al. in 2019, assessed hDPSCs functionality following sphere-mediated neural induction. In both cases, *in vitro* differentiated cells generated single AP-like fast depolarizations resembling the rising phase of neuronal APs. However, these cells failed to fully repolarize to baseline membrane potential and no more than one AP-like depolarization could be triggered on the same cell (9,10).

Recently, our group reported a new neurodifferentiation protocol, supplementing hDPSCs cultures with retinoic acid and pulses of 40mM potassium chloride, which promoted the generation of voltage-dependent potassium and sodium currents, spontaneous electrophysiological activity, as well as the triggering of repetitive neuronal APs with complete baseline recovery *in vitro* (12). Despite these advances, the ability of hDPSCs to form synaptic connections and integrate into functional neuronal networks remains largely unexplored. To date, only one *in vivo* study has examined the electrophysiological properties of hDPSCs following injection into the cerebrospinal fluid of newborn rats. Although most transplanted cells migrated to major brain neurogenic niches, they did not generate APs, and only voltage dependent sodium- and potassium-dependent currents were detected (16).

In the present study, we aimed to assess the capacity of hDPSC-derived cells to histologically integrate into the hippocampus of non-immunosuppressed C57BL/6J mice to evaluate whether these grafted cells could generate APs and establish functional synaptic interactions with the host tissue neurons. One-month after transplantation, GFP-transduced and grafted hDPSCs survived within the brain and exhibited an immature neuroblast-like marker phenotype characterized by doublecortin expression. Electrophysiological recordings revealed immature neuronal properties, such as high input resistances, low membrane capacitances, and the ability to fire only single APs, consistent with early stages of neuronal differentiation. Together, these findings expand current knowledge of hDPSCs electrophysiological behavior following *in vivo* brain integration and constitute a first step to investigate the temporal requirements for full neuronal maturation and functional incorporation of hDPSCs into host brain synaptic circuits.

## Materials and methods

### Isolation of hDPSCs and establishment of primary cultures

Primary cultures of hDPSCs were established as previously described (12,17). Briefly, human third molars from healthy donors were collected from dental surgery waste. Following tooth fracture, dental pulp tissue was carefully isolated and enzymatically digested for 1 h at 37°C using a solution containing 3 mg/mL collagenase (#17018029, Gibco) and 4 mg/mL dispase (#D4693-1G, Merck). After centrifugation, cell pellets from each donor were resuspended and plated onto tissue culture-treated flasks (#83.3912.002, Sarstedt) in DMEM (#D5796, Sigma-Aldrich) supplemented with 10% fetal bovine serum (FBS; #SV30160.03, HyClone), 100 U/mL penicillin and 150 mg/mL streptomycin (#11528876, Gibco). Cells were maintained under standard culture conditions to generate adherent monolayers cultures.

#### Lentiviral green fluorescent protein infection

For cell tracking, hDPSCs were transduced with lentiviral particles encoding green fluorescent protein (GFP), as previously reported (18). Monolayer cultures maintained in serum-containing DMEM media were used due to their higher proliferation rate and increased resistance to viral transduction. Cells were infected with the pLenti-CMV-GFP-Hygro vector (656–4) (Addgene viral prep # 17446-LV), kindly provided by Eric Campeu and Paul Kaufman, which confers resistance to hygromycin-B antibiotic (#31282-04-9, Thermo Fisher) and was purchased from the addgene non-profit repository (19).

To generate stable GFP^+^-hDPSCs populations, cells were seeded in 24 well-plates at a concentration of 5×10^4^ cells per well as previously reported (18). Briefly, after reaching confluence, lentiviral particles were added at a concentration of 2 x 10^5^ transduction units per milliliter (TU/ml). Twenty-four hours post infection, viral particles were removed and hygromycin-B (20 μg/mL; #31282-04-9, Thermo Fisher) was added to select stable transduced cells. GFP^+^-hDPSCs were subsequently expanded, and transduction efficiency was assessed by green fluorescence using a Nikon Eclipse Ts2 microscope.

#### Preparation of hDPSCs for intracerebral grafting

To obtain enough viable cells for intracerebral transplantation, a two-step expansion and neural priming protocol was applied. Following lentiviral transduction in serum-containing medium, GFP^+^-hDPSCs were cultured for 7 days in serum-free Neurocult proliferation to generate free-floating dentospheres and promote the expression of neural precursor markers prior to grafting (**Fig. 1a**).

**Figure 1:**
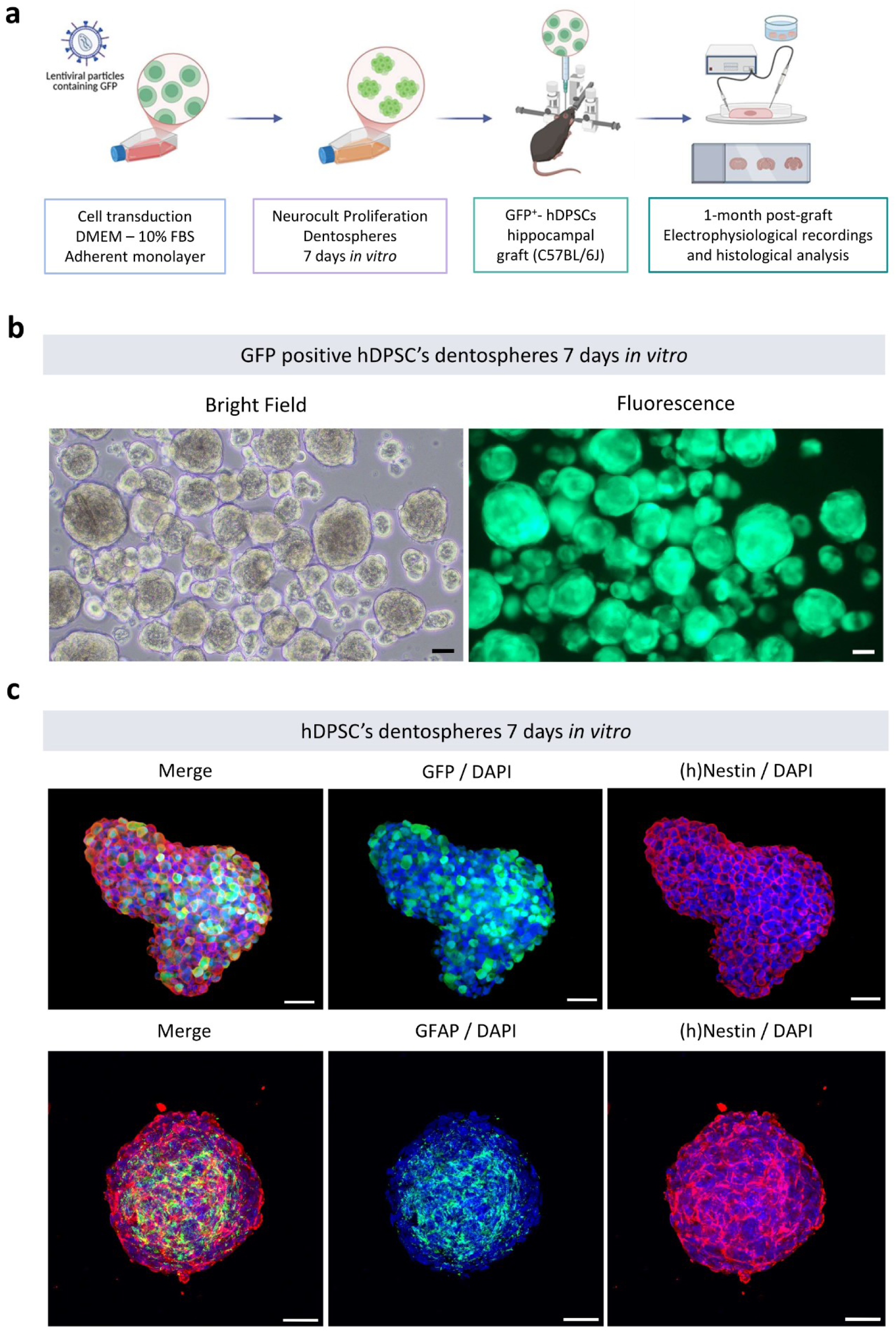
hDPSCs derived dentospheres neurogenic potential before their grafting into the mouse brain. **(A)** Schematic representation of the experimental procedure for cell transduction, dentospheres formation and hDPSCs grafting into the mouse brain (Created with BioRender). (**B**) Bright field and fluorescence images of GFP^+^-hDPSCs dentospheres transduced and grown in FBS containing media and thereafter transferred into serum-free proliferation media for 7 days. Scale bar: 100 µm. (**C**) Nestin and GFAP neural stem cell markers immune positive labeling in GFP positive dentospheres after 7 days into serum-free media. All the spheres are counterstained with DAPI. Scale bar: 50 μm.

Briefly, 5×10^5^ cells per well were seeded in low-attachment six well-plates (#83.3920.500, Sarstedt) containing Neurocult basal medium supplemented with human Neurocult proliferation supplement (9:1 ratio; #05751, Stem Cell Technologies), 2 μg/mL Heparin (#07980, Stem Cell Technologies), 20 ng/mL EGF and 10 ng/mL bFGF (#AF-100-15 / #AF-100-18B, Peprotech), 2% B27 without vitamin A (#12587010, Thermo Fisher), 2 mM L-Glutamine (#G7513, Sigma-Aldrich), 100 U/mL penicillin and 150 mg/mL streptomycin (#11528876, Gibco). Under these conditions, cells formed neurosphere-like dentospheres.

Immediately prior to transplantation, GFP expression within dentospheres was verified using an inverted fluorescence microscope (Nikon Eclipse Ts2). Culture supernatants were collected and centrifuged at 1,200 rpm for 10 minutes to recover the floating spheres. The resulting pellets were enzymatically dissociated with Accutase (#A6964-100ML, Sigma Aldrich) for 10 minutes at 37°C to obtain single-cell suspensions. Cell number, viability, and GFP transduction efficiency were subsequently quantified using a Luna-FL automated cell counter (#L20001, Logos biosystems), ensuring standardized and reproducible cell preparations for intracerebral injection.

#### Immunofluorescence of whole-mount spheres

Dentosphere characterization was performed by whole-mount immunocytochemistry on free-floating spheres. Briefly, collected spheres were centrifuged at 1,200 rpm for 10 minutes and the resulting pellet was fixed without disaggregation in 4% paraformaldehyde (PFA, #158127-500G, Sigma-Aldrich) overnight at 4 °C to preserve the three-dimensional architecture. Samples were then washed three times in phosphate-buffered saline (PBS; #D5652, Sigma-Aldrich) for 10 minutes and permeabilized overnight at 4°C in PBS containing 2% Triton X-100 (#93443, Sigma-Aldrich). All the steps of this protocol were performed under agitation, with centrifugation at 1,200 rpm for 10 minutes between steps.

Prior to antibody incubation, spheres were washed three times during 10 minutes with PBS and blocked overnight at 4°C in PBS supplemented with 1% Triton X-100 and 1% bovine serum-albumin (BSA; #A9647, Sigma-Aldrich). Neural stem cell marker expression was assessed using anti-human Nestin (1:200; #MAB1259, Biotechne R&D) and anti-Glial Fibrillary Acidic Protein (GFAP; 1:400; #G9269, Sigma-Aldrich) diluted in PBS containing 1% Triton X-100 and 0.5% BSA. Spheres were incubated with primary antibodies for 72 h at 4 °C. Following incubation, samples were washed three times for 30 minutes in PBS.

Secondary antibodies (Alexa Fluor 488 and Alexa Fluor 555 donkey anti-rabbit and anti-mouse, respectively (1:250; #A21206, #A31570, Invitrogen) together with DAPI (1:500; #10116287, Thermo Fisher Scientific) were applied overnight at 4°C in PBS containing 1% Triton X-100 and 0.5% BSA. After three final washes in PBS (30 minutes each), spheres were mounted as a single drop on glass slides. Immunofluorescence images were acquired a LSM800 confocal microscope (Zeiss, Jena, Germany).

#### hDPSCs grafts into the mouse brain

Animal procedures were conducted in compliance with Council Directive 90/219/EC (23 April 1990), as amended by Directive 2005/174/EC (05 March 2005), and with European Union legislation governing the protection of animals used for scientific purposes (Directives 86/609/EEC & 2010/63/EU). Experimental protocols were approved by the Ethics Committee of the University of the Basque Country (EHU) and the competent authority of the Provincial Administration of Bizkaia for the use of human cells and experimental animals (CEISH M10/2023/025; CEEA M20/2023/026 and M30/2023/027), as well as by the ethics committees of the Interdisciplinary Institute of Neuroscience in Bordeaux and the relevant French authorities (project reference: 46334, dossier reference: 202312181625835). Animals were housed in colony isolators under 12 h light/dark cycles with food and water *ad libitum*.

One hour prior to transplantation, GFP^+^-hDPSC dentospheres were enzymatically dissociated using Accutase and resuspended in Neurocult serum-free basal medium, as previously described (20). Two-months-old C57BL/6J mice were used as hosts for hDPSCs *in vivo* grafting. Anesthesia was induced with 4% isoflurane in a closed chamber at an oxygen flow rate of 1.5 L/min and maintained throughout surgery at 2% isoflurane via facemask with an oxygen flow of 0.4 L/min. Mice were secured in a stereotaxic apparatus (Kopf model 900) using ear bars positioned in the external auditory meatus. Animal head was shaved and disinfected with 10% betadine solution.

To provide pre-emptive analgesia, mice received a local injection of Lidocaine (100 μl, 7 mg/kg) and a subcutaneous injection of buprenorphine (300 μl, 0.1 mg/kg) 5 minutes before surgery. During the procedure, corneal dehydration was prevented by application of ophthalmic ointment (Vaseline), and body temperature was maintained using a heating pad. With the help of a scalpel, a midline incision was made cutting the skin and exposing the skull. Bregma cranial suture was used as stereotaxic reference point. A small burr hole was drilled using a micro drill (# 51449, Stoelting, Illinois, USA).

Cell suspension (2 μl containing 50,000 cells) was injected bilaterally into the hippocampus at a rate of 0.5 μl/min using a micropump attached to a 10 μl Hamilton syringe with a 33G needle (Hamilton, Bonaduz, Switzerland). Injection coordinates relative to bregma were: Anteroposterior (AP): −2 mm, laterolateral (L): +/- 1.5 mm and dorsoventral (DV): 1.5 mm. Pre- and post-operative care was performed according to established protocols previously described (20).

A total of 12 mice (7 males and 5 females) received GFP^+^-hDPSCs grafts. One month post graft, three animals were perfused for histological analysis while the remaining animals were used for electrophysiological recordings in acute brain slices.

#### Electrophysiological recordings in acute brain slices

One month after grafting, animals were anesthetized with 5% isoflurane for 2 minutes and immediately decapitated. Brains were rapidly removed and transferred to the chamber of a vibratome (Leica VT1200s) containing carbogen (95% O_2_, 5% CO_2_) and an ice-cold cutting solution composed of (in mM): 2 KCl, 2.6 NaHCO_3_, 1.15 NaH_2_PO_4_, 10 D-Glucose, 120 Sucrose, 0.2 CaCl_2_ and 6 MgCl_2_. Serial hippocampal sections (350 μm) were cut at a speed of 0.05 mm/s and incubated during 30 minutes at 35 °C in artificial cerebrospinal fluid (aCSF) containing (in mM): 125 NaCl, 2.5 KCl, 1 MgCl_2_, 1.25 NaH_2_PO_4_, 26 NaHCO_3_, 16 D-Glucose and 2 CaCl_2_ (pH = 7.3). The osmolarity of aCSF was checked with the osmometer and adjusted to 290-310 mOsm/L. This solution was used both for slice recovery and as extracellular recording solution, continuously bubbled with carbogen (95% O_2_, 5% CO_2_) throughout the experiments to preserve tissue viability.

Whole-cell current-clamp recordings of grafted GFP^+^-hDPSCs were performed at 30-32 °C using a fluorescence microscope (Olympus TH4-200, Olympus Optical, Japan) provided with a 40x water immersion lens and MultiClamp 700B amplifier (Molecular Devices, San Jose, CA). Signals were filtered at 2.9 kHz and acquired at a 10 kHz sampling rate using a DigiData 1550 data acquisition system and pCLAMP 10.3 software (Molecular Devices, San Jose, CA). Patch clamp pipettes were made of borosilicate glass (#30-0053, Warner Instruments) and had a resistance of 3–5 MΩ with an internal solution containing (in mM): 120 K-gluconate, 20 K-chloride, 10 HEPES, 10 phosphocreatine, 4 Mg-ATP, and 298 Na-GTP (292 mOsm, pH = 7.4). Fast and slow whole-cell capacitances were neutralized. In current-clamp mode, action potentials (APs) were evoked by step of positive current injections lasting 200 ms. This protocol was repeated 33 times, with the injected current increasing by 25 pA with each iteration, starting at −200 pA and reaching up to 600 pA. An interval of 400 ms was applied to allow the recovery of the cells between stimuli. In selected recordings, tetrodotoxin (TTX, 1 μM; #554412, Sigma Aldrich) was used to block the evoked APs. Obtained data were analysed offline using Igor Pro 8 software (Wave Metrics).

#### Animal perfusion and brain processing

One month after hDPSCs transplantation, animals were deeply anaesthetized by intraperitoneal injection of ketamine (100 mg/kg) and xylazine (20 mg/kg) and intracardially perfused with first phosphate-buffered saline (PBS) and 4% paraformaldehyde (PFA). Brains were carefully removed and sectioned using a vibratome (Leica VT1200s) into serial coronal sections of 50 μm thickness at a cutting speed of 6 mm/s and a frequency of 6 Hz. Free-floating sections were collected and stored in PBS containing 0.2 % sodium azide (#S2002, Sigma-Aldrich) at 4 °C until further histological processing.

#### Immunohistochemistry in brain slices

Immunofluorescent staining was performed on free-floating sections following a standard protocol with minor adaptations (21). Briefly, sections were permeabilized in PBS containing 1.5% Triton X-100 for 30 minutes at room temperature, followed by two washes of 5 minutes in PBS with 0.1% Triton X-100. Non-specific binding was blocked by incubating the sections for 1 h in PBS containing 0.1% Triton X-100 and 5% bovine serum albumin (BSA). Primary antibodies were incubated overnight in PBS solution containing 0.3% Triton X-100, 2% BSA. Grafted human cells were identified using an anti-GFP antibody (1:1000; #ab315212, Abcam), anti-STEM121 (1:1000; #Y40410, Takara) and anti-human Nestin (1:200; #MAB1259, Biotechne R&D). Stemness was assessed using anti-SOX2 (1:500; #ab97959, Abcam). Immature neuroblasts were detected with anti-DCX (1:300; #F6K9E, Cell Signalling Technology), while mature neurons were identified using antibodies against β-III-Tubulin (1:250; #ab18207, Abcam) and NeuN 1:500 (#ab177487, Abcam). Astrocytes and microglia were labelled with anti-GFAP (1:400; #G9269, Sigma-Aldrich) and anti-Iba1 (1:1000; #ab289874, Abcam) respectively.

After primary antibody incubation, sections were washed three times for 10 minutes in PBS containing 0.1% Triton X-100 and then incubated for 2 h at room temperature with appropriate donkey secondary antibodies conjugated to Alexa Fluor 488, 555, or 647 (1:500; #A31572, #A32766, #A21432, #A32795 Invitrogen), together with DAPI (1:1000; #10116287, Thermo Fisher) for nuclear counterstaining. Sections were subsequently washed three times in PBS and mounted on glass slices.

Fluorescent images were acquired using a Zeiss LSM800 confocal microscope and a Zeiss Apotome 2 fluorescence microscope (Jena, Germany) equipped with 20x and 40x objectives. Whole-brain serial sections were digitized using a 3D Histech Slide Scanner (Budapest, Hungary).

#### Quantitative image analysis

To analyze the brain-wide distribution of GFP^+^-hDPSCs, whole brains were serially sectioned, mounted, and digitized as tiled images using 3D Histech Slide Scanner. Images were analyzed manually with Image-J public software (version 1.50e) (22) and automatically using the QUINT workflow, which enables spatial registration of histological sections to the Allen Mouse Brain Atlas for accurate anatomical localization of the transplanted cells. Briefly, images were first converted into a compatible format using Nutil. Coronal sections were then automatically registered to the reference atlas using DeepSlice (23,24), followed by manual refinement in QuickNII and Visualign to ensure anatomical precision. GFP-positive pixels were subsequently identified by supervised classification with Ilastik, quantitative features were extracted with Nutil, and three-dimensional reconstruction was generated using MeshView.

Co-localization of GFP^+^-hDPSCs with neuronal markers was assessed using the Just Another Colocalization Plugin (JACoP) for Image-J (25). Images were acquired from the hippocampus of both hemispheres, from two coronal sections per mouse, in a total of three mice. Co-localization was quantified by calculating Mander’s coefficient, which measures the fraction of signal overlap between two channels, and Pearson’s correlation coefficient (R), which assesses the linear correlation of fluorescence intensity values between the two channels.

### Statistical analysis

Data were analyzed using GraphPad Prism 8 software (Boston, MA, USA). According to the characteristics of the data distribution, comparisons between two groups were made via either Student’s t-test or the Mann‒Whitney U test. P values less than 0.05 were considered statistically significant. The results are presented as mean ± SD.

## Results

### Lentiviral GFP transduction of hDPSCs resulted in dentospheres that stably expressed GFP and neural stem cell markers

Cell labeling through viral transduction, which enables stable genomic integration of fluorescent reporter proteins, is essential for the identification of live grafted cells within host tissues. This strategy allows real-time visualization of grafted human cells by fluorescence microscopy during live recordings (19,26,27). Although neural commitment of hDPSCs is optimally achieved under serum-free medium, this approach markedly decreases cell harvesting yield (28,29). In contrast, efficient lentiviral transduction requires a high density of healthy, proliferating cells. To reconcile these opposing requirements, we implemented a two-phase protocol in which hDPSCs were initially expanded and transduced in FBS containing medium, followed by a switch to serum-free proliferation medium to enhance neurogenic potential prior to brain grafting (**Fig. 1a**). Primary hDPSC cultures were efficiently transduced *in vitro* with lentiviral particles encoding GFP reporter gene and Hygromycin-B resistance. GFP expression remained stable across passages under both proliferation conditions and transduced cells retained their capacity to form dentospheres despite transient serum exposure during transduction (**Fig. 1b**). Importantly, these dentospheres expressed the neural markers Nestin and GFAP, confirming neurogenic potential before transplantation into the mouse brain (**Fig. 1c**).

### GFP positive hDPSCs were found in C57BL/6J mouse hippocampus one month after their graft into the CA1 region

One of the principal challenges of stem cell grafting into the brain is the risk of immune rejection by the host tissue (30). Following transplantation, microglia, macrophages and lymphocytes typically accumulate around the graft site and often display an activated phenotype (31). After transplantation of GFP positive neural-preconditioned hDPSCs into C57BL/6J mice, grafted cells were readily identified by fluorescence microscopy within the injected brain region (**Fig. 2a**). One-month post-grafting, GFP^+^-hDPSCs were predominantly localized into the CA1 region of the hippocampus, corresponding to the primary injection site (**Fig. 2b-b.1**), but were also detected in adjacent regions, including the dentate gyrus (**Fig. 2b.2**) and the corpus callosum (**Fig. 2c**). Notably, despite the absence of immunosuppression, GFP^+^-hDPSCs survived within the C57BL/6J brain, and in regions distal to the injection site the surrounding microglia and astrocytes did not exhibit overtly reactive morphology or evidence of GFP^+^ particle internalization (**Fig. 2d**).

**Figure 2:**
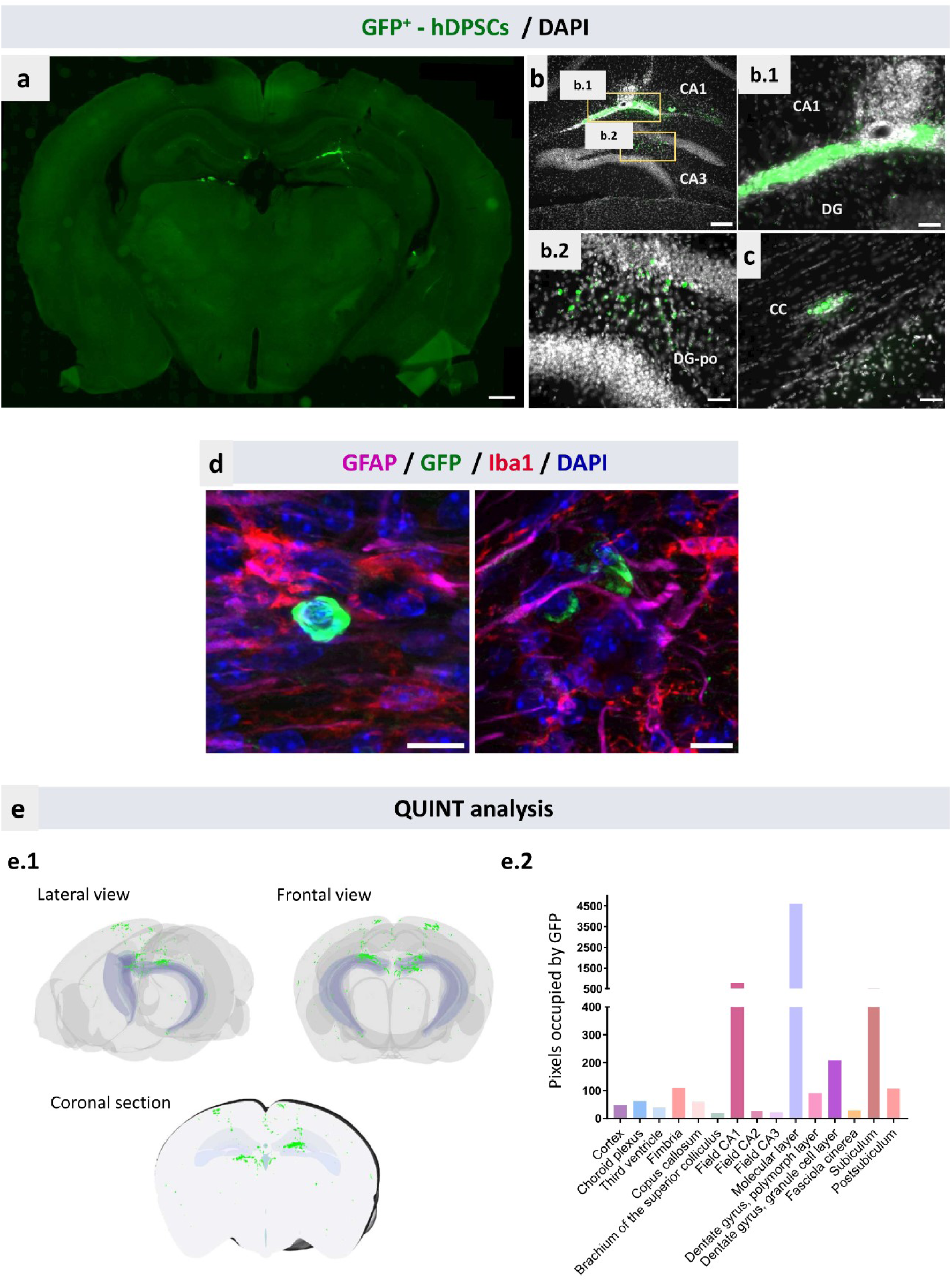
GFP^+^-hDPSCs into C57BL/6J mouse brain at one month after hippocampal transplantation. **(A)** Coronal section showing the area where GFP^+^-hDPSCs (green) were injected one month after grafting them. **(B)** Hippocampal CA1 and CA3 regions showing a positive GFP immune labeling above the dentate gyrus (DG) in the molecular layer **(B.1)** and into the dentate gyrus polymorph layer (DG-po) **(B.2)**. **(C)** GFP positive labeling in the corpus callosum (CC). All the images are counterstained with DAPI. Scale bar 50 μm. **(D)** GFP^+^-hDPSCs between microglial Iba1 positive cells (red) and GFAP labelled astrocytes (magenta) in the CA1 region. Scale bar of 10 μm. (**E**) Grafted cells dispersion results measured by the QUINT bioinformatics workflow. (**E.1**) Three-dimensional reconstruction of an entire grafted brain obtained by the Mesh View Atlas Viewer and (**E.2**) graph showing the number of pixels occupied by GFP positive cells in each brain region measured by Nutil software.

The spatial dispersion and precise localization of grafted cells were quantitatively assessed using QUINT bioinformatics workflow, which aligns GFP^+^-hDPSCs positive pixel signal with the Allen Mouse Brain Atlas. Three-dimensional reconstruction of the entire grafted brain one-month post-transplantation revealed that GFP expression was predominantly distributed bilaterally within the hippocampus and surrounding areas (**Fig. 2e.1**). Quantitative analysis showed that the highest GFP signal was detected in the CA1 region (801 pixels) and the CA1 molecular layer (4610 pixels), corresponding to the grafting sites (**Fig. 2e.2 and Fig. 2b**). In addition, GFP^+^-hDPSCs were detected in key adult neurogenic regions, including the third ventricle (39 pixels) and the dentate gyrus granular (209 pixels) and polymorph (90 pixels) layers (**Fig. 2e.2**). Outside the hippocampus, smaller GFP^+^ signals were observed in the cortex (47 pixels) and corpus callosum (60.3 pixels), indicating a limited but measurable cell migration beyond the primary graft area (**Fig. 2e.2 and Fig. 2c**).

Even when immune barriers are effectively overcome by grafted cells, their post-transplantation identification remains challenging. Several studies have reported a progressive downregulation of GFP expression in grafted cells *in vivo* (32–34). Consistent with these observations, quantitative analysis of GFP^+^/DAPI cells in coronal sections of an entire injected brain using Image J revealed that, despite the implantation of 100,000 cells per brain, only 2,751 GFP^+^-hDPSCs were detectable one month after grafting (**Fig. 3a**). This finding suggests that a proportion of transplanted cells either downregulated GFP expression or were lost during or after the transplantation process. Therefore, to further determine whether grafted cells had or not persisted despite a possible GFP silencing, we performed immunolabeling with STEM121, a human-specific marker widely used to detect the engraftment, migration and differentiation of transplanted human cells in murine tissues (35). Although a subset of cells showed co-expression of GFP and STEM121, a substantial number of STEM121^+^ cells lacked detectable GFP expression (**Fig. 3b**), indicating that GFP downregulation rather than complete cell loss contributed more significantly to the underestimation of graft survival based solely on GFP labeling.

**Figure 3:**
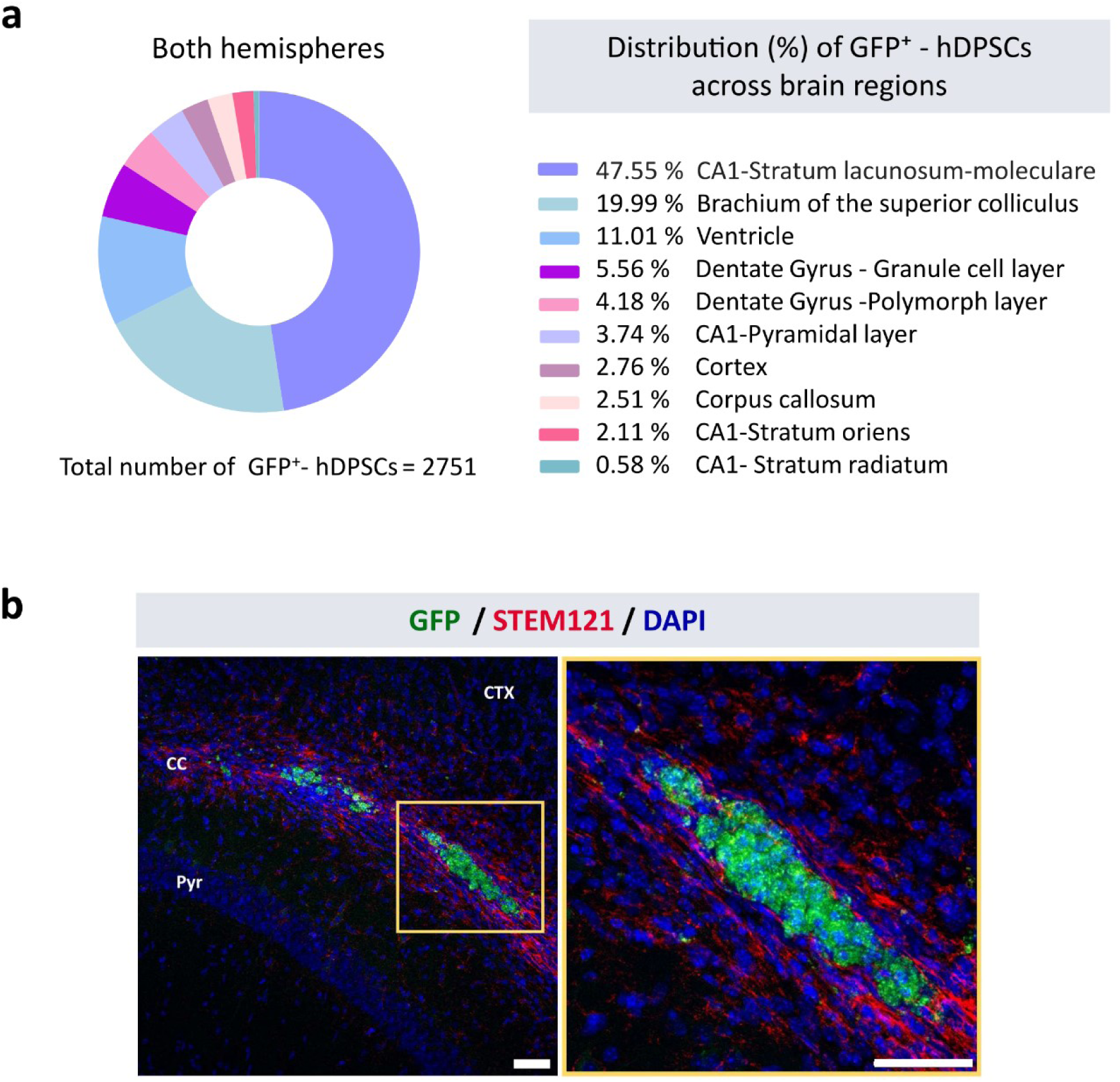
STEM 121 expression in hippocampal grafted hDPSCs. **(A)** Percentage of GFP^+^-hDPSCs per region quantified in both hemispheres with Image J software. **(B)** Confocal images showing hDPSCs labeling with STEM 121 (red) and GFP (green) markers in the hippocampus. Scale bar of 50 µm.

### GFP^+^-hDPSCs expressed immature neuronal markers one month after grafting into the brains of C57BL/6J mice

During physiological neural differentiation in the brain, neural stem cells progress through distinct maturation stages characterized by the expression of specific molecular markers (**Fig. 4a**) (36). One-month post transplantation, only a small fraction of hDPSCs retained features of an undifferentiated state. Sparse GFP^+^-hDPSCs located in the CA1 hippocampal region and cortical areas surrounding the injection site exhibited positive immunolabeling for the human cytoskeletal stem cell marker Nestin (**Fig. 4b**). In addition, Sox2, a transcription factor related with the stemness and pluripotency of the cells, was not detected in GFP^+^-hDPSCs (**Fig. 4c**).

**Figure 4:**
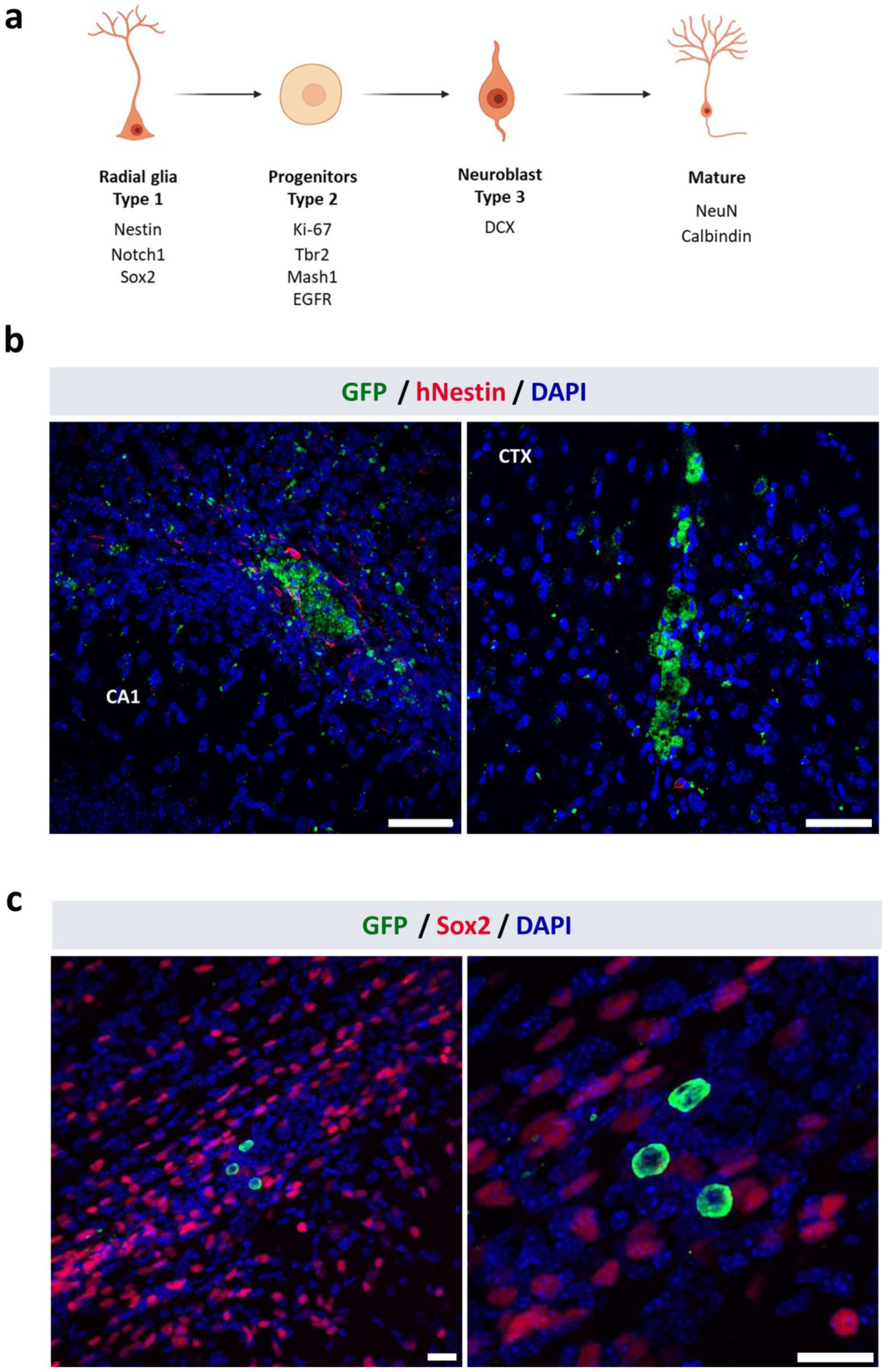
Human Nestin and Sox2 positive labeling in C57BL/6J mouse brain at one month post transplantation. **(A)** Scheme showing the different states and markers of neural stem cells during the neurodifferentiation process (Created with BioRender). (**B**) hDPSCs labelled with an antibody against human Nestin (red) and GFP in CA1 region and into the cortex (CTX). Scale bar 50 μm. (**C**) Sox2 nuclear staining (red) in GFP^+^- hDPSCs (green) into the hippocampal neurogenic area. Scale bar 20 μm.

Notably, expression of the immature neuronal marker doublecortin (DCX) was primarily observed in two hippocampal regions. Endogenous mouse neuroblasts were detected within the dentate gyrus neurogenic niche, whereas GFP^+^/DCX^+^ hDPSCs were identified in the CA1 region adjacent to the corpus callosum (**Fig. 5a**). Co-localization analysis using the Just Another Colocalization Plugin (JACoP) for Image-J revealed that 62.8% of the DCX signal overlapped with GFP fluorescence (**Fig. 5b**) while of 65.9% of GFP^+^-hDPSCs were located within DCX-positive areas (**Fig. 5c**). Pearson’s correlation coefficient, which assesses pixel-by-pixel intensity correlation between fluorescence channels, indicated a moderate positive correlation (R = 0.5) between DCX and GFP signals in the grafted brains (**Fig. 5d**).

**Figure 5:**
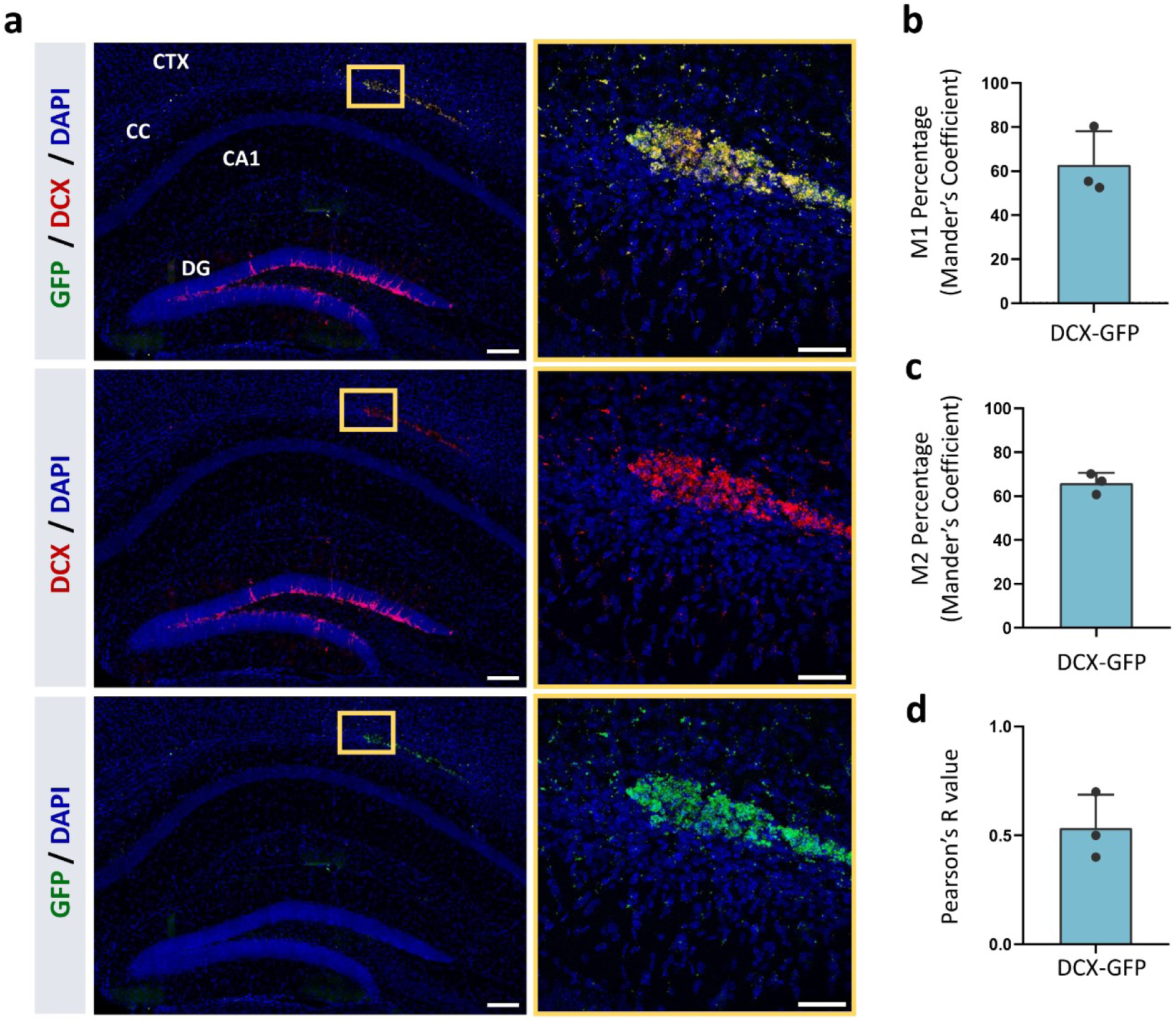
Doublecortin (DCX) labeling into the hippocampus of GFP^+^-hDPSCs injected mice at one month post graft. **(A)** Confocal images showing DCX positive signal in red and hDPSCs labeled with GFP in green and the merge signal resulting from the co-localization of both channels. Scale bar for the entire hippocampus tiles images of 20 μm and for the inset 50 μm. **(B)** Mander’s coefficient 1 showing the intensity amount from red channel (DCX) that co-localizes with green channel (GFP). **(C)** Mander’s coefficient 2 showing the amount of intensity from green channel (GFP) that co-localizes with red channel (DCX). **(D)** Pearson’s R value showing the linear relationship between the intensities of red and green channels pixel by pixel. (n=3 C57BL/6J; CTX: Cortex, CC: Corpus callosum and DG: Dentate gyrus).

To further evaluate neuronal maturation, the expression of more mature neuronal markers and complex neuronal morphologies were examined in the hippocampus. GFP^+^ grafted cells did not exhibit immunoreactivity for NeuN mature neuronal marker (**Fig. 6a)**, nor did they display a characteristic βIII-tubulin-positive axo-dendritic neuronal phenotype (**Fig. 6b**). Co-localization analysis showed that only 0.75% of the βIII-tubulin signal overlapped with GFP (**Fig. 6c**). However, Mander’s second coefficient revealed that 28.24 % of the GFP^+^-hDPSCs were located in regions containing βIII-tubulin positive structures, but without a direct co-localization (**Fig. 6d**). Consistently, Pearson’s correlation coefficient was low (R = 0.11), further supporting the absence of a significant association between grafted GFP^+^ cells and the expression of mature axo-dendritic neuronal markers (**Fig. 6e**).

**Figure 6:**
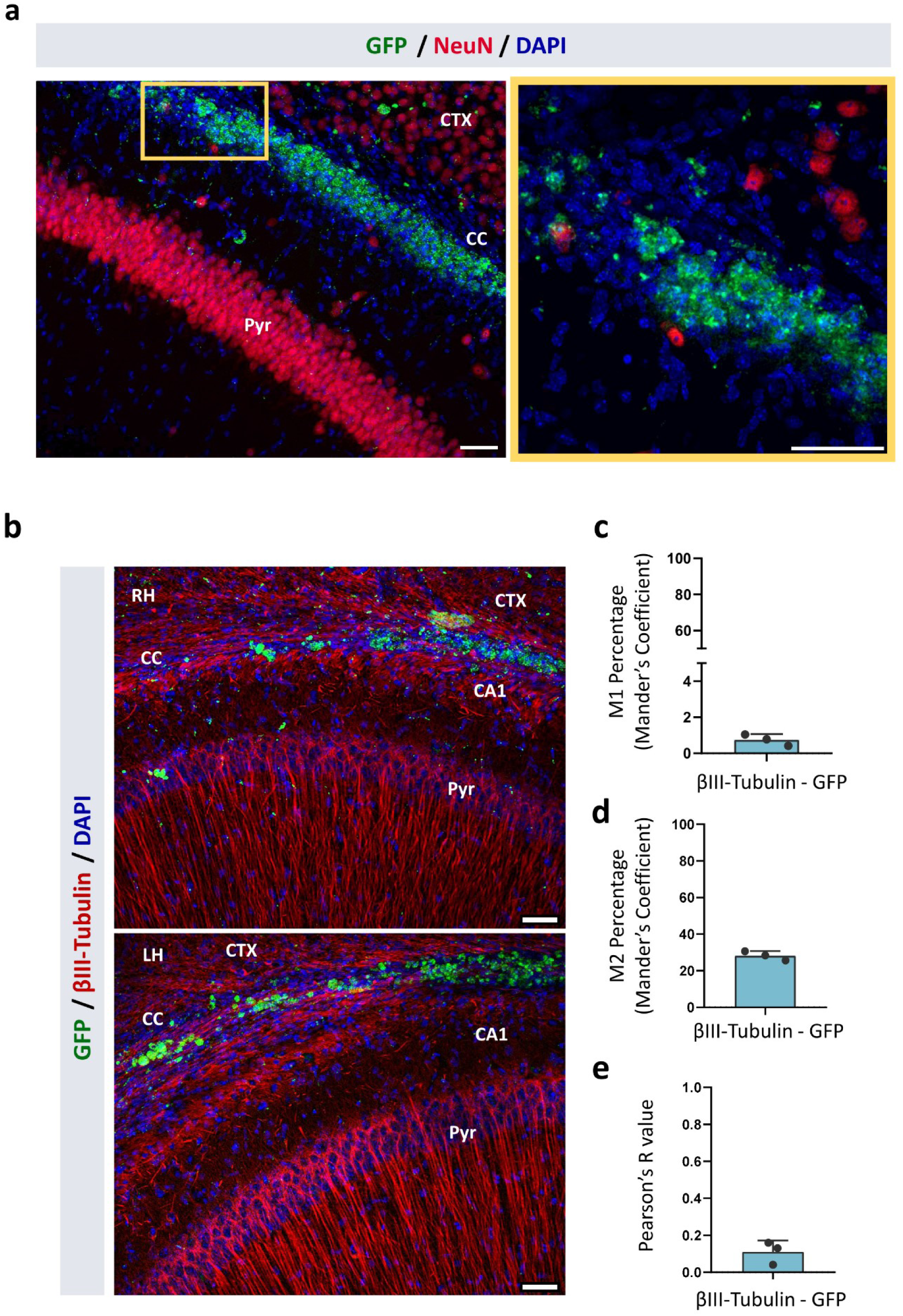
GFP, NeuN and βIII-Tubulin labeling into the hippocampus of grafted mice at one month post transplantation. **(A)** Confocal images showing NeuN positive signal in red and hDPSCs labelled with GFP. Scale bar of 50 µm. **(B)** Confocal images showing neuronal fibre tracks labeled in red with βIII-Tubulin and GFP^+^-hDPSCs in green into the right hemisphere (RH) and the left hemisphere (LH). Scale bar 50 μm. (CTX: Cortex, CC: Corpus callosum and Pyr: CA1-Pyramidal layer). **(C)** Mander’s coefficient 1 showing the amount of intensity from red channel (βIII-Tubulin) that co-localizes with green channel (GFP). **(D)** Mander’s coefficient 2 showing the amount of intensity from green channel (GFP) that co-localizes with red channel (βIII-Tubulin). **(E)** Pearson’s R-value showing the linear relationship between the intensities of βIII-Tubulin and GFP pixel by pixel (n=3).

### GFP^+^-hDPSCs displayed immature, neuroblastic-like electrophysiological properties one month after injection into the hippocampus of C57BL/6J mice

Immature neuroblasts exhibit electrophysiological properties different from those of mature neurons, including higher input resistances and more depolarized resting membrane potentials (37,38). To characterize the electrophysiological profile of GFP^+^-hDPSCs that displayed a DCX-positive, neuroblast-like phenotype in histological analyses, whole-cell patch-clamp recordings were performed in acute hippocampal brain slices from grafted C57BL/6J mice. One month after graft, small, round GFP^+^-hDPSCs were identified by green fluorescence within the hippocampus and a total of 11 GFP^+^ cells from four different animals were recorded (**Fig. 7a**).

**Figure 7:**
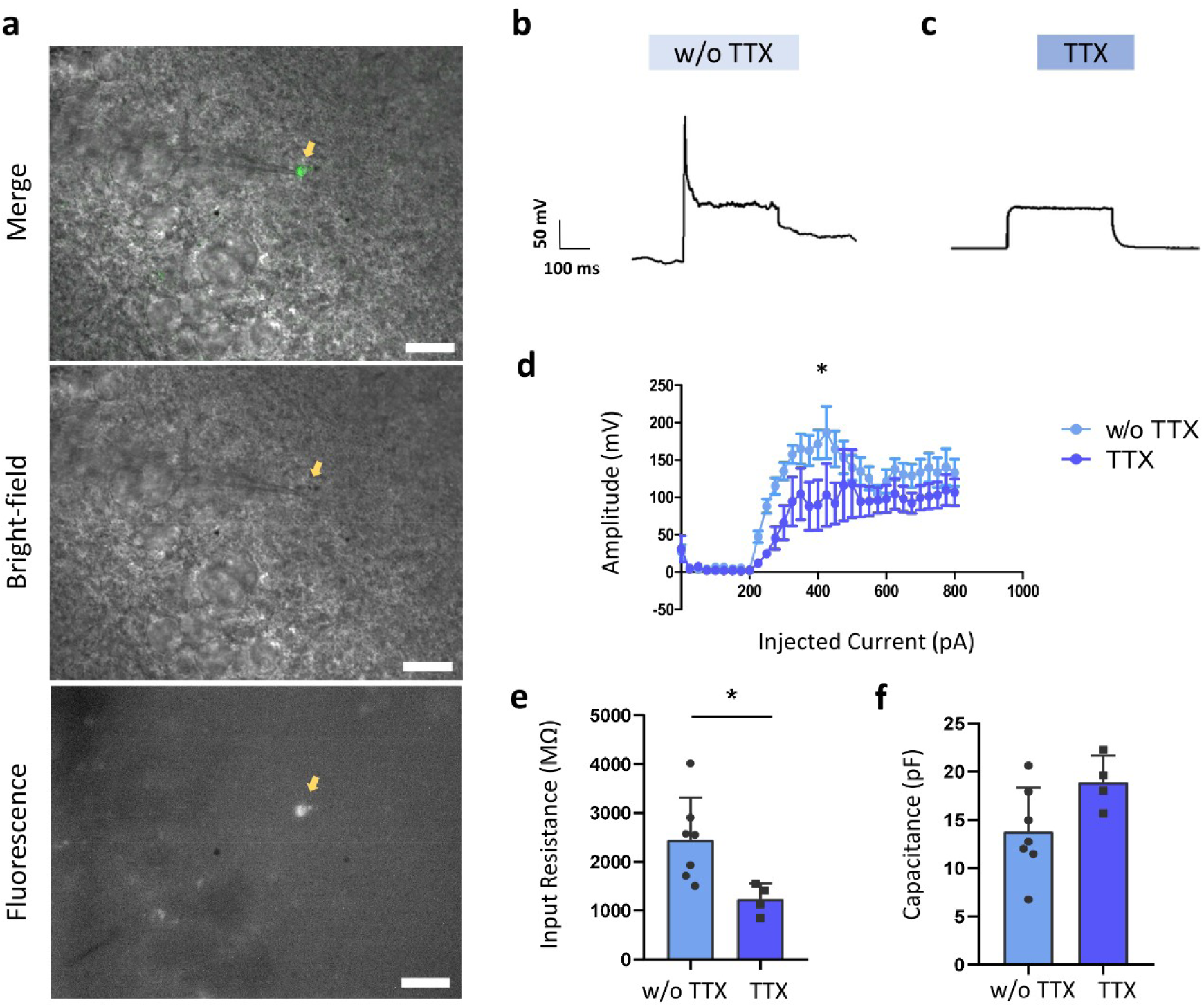
*Ex vivo* electrophysiological recordings from grafted GFP^+^- hDPSCs at one month post transplantation into the mouse hippocampus. **(A)** Bright-field and fluorescence images showing the appearance of the recorded GFP positive cells. **(B)** Single APs elicited by depolarizing GFP^+^-hDPSCs. **(C)** Blockade of the APs when 1 μM tetrodotoxin (TTX) was perfused into the recording chamber**. (D)** Graph showing the mean amplitude (mV) of the registered single APs with and without TTX along the growing range of injected current (pA). **(E)** Input resistance mean values in mega ohms (MΩ) for GFP positive recorded cells with and without TTX blockade. **(F)** Capacitance mean values in picofarad (pF) for registered cells with and without TTX. * p < 0.05 Student t-test or Mann‒Whitney U test (Two-tailed).

In current clamp mode, cells were depolarized with 33 consecutive current injections (25 pA increments; 200 ms duration). Depolarizing current injections consistently evoked single APs however, repetitive firing was not observed (**Fig.7b**). Bath application of 1 μM TTX completely abolished these APs, demonstrating that spike generation was mediated by neuronal voltage-gated sodium channels (**Fig. 7c**). Evoked APs exhibited a mean peak amplitude of 178.155 ± 30.83 mV, which was significantly reduced to 98.67 ± 42.54 mV following TTX perfusion (p = 0.0217, **Fig. 7d**).

Recorded GFP^+^-hDPSCs presented high mean input resistance values (2457.39 MΩ; **Fig. 7e**) and low membrane capacitances (13.79 pF; **Fig. 7f**), both electrophysiological features of immature neuronal cells. Notably, upon TTX application, input resistance values were significantly reduced to 1319.52 MΩ (p = 0.0244), supporting a possible contribution of sodium channel activity to the basal membrane properties of grafted cells (**Fig. 7e**).

## Discussion

After a traumatic, degenerative or inflammatory process, the regenerative potential of the human brain is very limited, and the currently available treatments only relieve symptoms without restoring the damaged tissue. Therefore, stem cell transplantation into the central nervous system (CNS) has become a major research field for replacing damaged or lost neurons in these pathologic contexts (39,40). However, the suitability of the stem cell grafts for restoring neuronal connectivity critically depends on the capacity of the donor cells to *in situ* differentiate to mature viable neurons that engage in synaptic interaction within the host brain circuitry. In addition, appropriate models are required to assess this neuronal integration process into a relevant preclinical scenario. While transplantation represents a key tool for assessing the survival and *in vivo* capability of stem cells for neural differentiation and replacement, very little is known yet about the functional integration of hDPSCs into the host brain circuitry (16). Our study demonstrates that neural-preconditioned, GFP-labelled hDPSCs can survive, engraft, and disperse within the mouse hippocampus in the absence of immunosuppression, while maintaining detectable human-specific identity despite partial GFP silencing. Importantly, grafted cells adopted an immature neuronal-like phenotype, as evidenced by DCX expression and neuroblast-like electrophysiological properties, yet failed to achieve full neuronal maturation, lacking expression of mature neuronal markers and functional integration within host circuits. Collectively, these findings highlight both the *in vivo* neurogenic potential and the intrinsic limitation of hDPSCs to reach complete neuronal differentiation within one month after transplantation.

Although hDPSCs capacity to survive and interact with CNS and peripheral nervous system (PNS) cells after transplantation has been widely demonstrated in different animal models of neurodegeneration and traumatic brain injury (41–47), their electrophysiological properties have been only described in a limited number of studies *in vitro*. Gervois and Li were the first to report action potential-like fast depolarizations in neurodifferentiated hDPSCs (9,10). More recently, our research group developed an improved neurodifferentiation protocol in which retinoic acid and potassium chloride treated hDPSCs displayed voltage-dependent potassium and TTX sensitive sodium currents as well as spontaneous electrophysiological activity and repetitive neuronal APs with a full baseline potential recovery (12). Nevertheless, hDPSCs functionality and capacity to integrate into the synaptic network *in vivo* has not yet been proved. To the best of our knowledge, the present study provides the first evidence of hDPSCs neural differentiation after their graft into immunosuppressed mouse hippocampus giving rise to immature neuronal like cells displaying APs firing activity.

To unequivocally demonstrate that the engrafted stem cells have survived, integrated and differentiated into the host tissue, cell labeling is a crucial step prior to transplantation (48). In xenotransplantation models, such as human-to-mouse grafts, human-specific antibodies including human nuclei (HuNu) (49,50), STEM 121 (51,52) or human Nestin (53,54) are commonly employed for post-mortem identification of grafted human cells in fixed tissues. However, for functional studies such as electrophysiological recordings in acute brain slices or multiphoton optogenetics in freely behaving animals, these immunohistochemical markers are insufficient because they require tissue fixation. To identify live grafted cells or to explore host-to-graft synaptic connections in live tissues, viral transduction allowing stable integration of fluorescent reporter proteins prior to transplantation is required. This approach enables real-time visualization of grafted human cells during live recordings (26,27,55). Therefore, in this study, we generated a stable GFP^+^-hDPSCs population using lentiviral particles carrying GFP, allowing localization of grafted cells in viable tissues for functional characterization.

Successful lentiviral transduction strongly depends on optimizing several culture parameters to achieve high transduction efficiency while preserving cells viability. Target cells must be at an appropriate density and in physiologically healthy state (56). Even though hDPSCs tend to acquire neural-like phenotypes under serum-free conditions (12), removal of FBS from the culture medium substantially reduces cell expansion rates, despite not significantly affecting cell viability (54,57). For this reason, several studies have established two-phase expansion protocols, in which cells are initially expanded with serum-containing media to reach the required cell number and subsequently transferred to serum-free conditions to enhance their neurogenic potential (9,10). Prior to GFP^+^-hDPSCs grafting into the mouse hippocampus, we adopted a similar two-phase proliferation protocol. Lentiviral transduction efficiency was optimized under serum-containing conditions, followed by promotion of a neurogenic phenotype through dentosphere formation.

Successful integration of the grafted cells within the host tissue, which can release immunomodulatory or neurotrophic factors that contribute to functional recovery, is strictly dependent on the number of surviving cells, as only viable cells can exert paracrine effects and replace damage tissue (58). Therefore, to enhance cell survival and migration to the target location after grafting, it is essential to carefully select the timing, number of cells, and injection site (59,60). In our study, we grafted GFP^+^- hDPSCs into the CA1 region of the hippocampus, an area characterized by a high plasticity which, under physiological conditions, integrates new-born neurons into its synaptic network (61). Moreover, graft cell density plays a critical role in survival: evenly distributed cells benefit from improved access to oxygen and nutrients, whereas densely packed grafts suffer from nutrient deprivation, promoting apoptosis within the first 24 hours after injection (58,60). Based on these considerations, we injected 50,000 cells per hemisphere. Regarding timing, immature neurons without extensive axonal projections exhibit improved survival and growing rates; therefore, cells should be harvested near cell-cycle exit, after genetic neuronal commitment but prior to extensive process outgrowth (40,62).

Beyond grafting parameters, activation of the immune system and subsequent rejection of transplanted cells represent another mayor challenge. Graft survival depends largely on innate immune responses and the local chemical microenvironment generated during transplantation. Even under physiological healthy conditions, cell grafting into the brain triggers an acute inflammatory response, with microglia and astrocytes rapidly colonizing the grafting site. This pro-inflammatory milieu can compromise graft survival and fate (31,58,63). Nevertheless, hDPSCs have been shown to modulate microglial reactivity, both *in vitro* by reducing LPS-induced pro-inflammatory cytokine expression and *in vivo -* by decreasing the number of hyperreactive microglia in the hippocampus of Alzheimer’s disease mouse models (64). In our hands, despite grafting into non-immunosuppressed mice, GFP^+^-hDPSCs survived in the brains of C57BL/6J mice. Moreover, in regions distant from the injection site, surrounding microglia and astrocytes did not show evidence of GFP^+^ particle internalization, as reported in previous GFP^+^ cell grafts studies (65).

Under proliferative conditions *in vitro*, neural precursors maintain stable GFP expression without signs of toxicity or transgene silencing. In contrast, *in vivo* differentiation and migration of grafted stem cells has been associated with downregulation of GFP expression, suggesting a close relationship between neural differentiation and transgene silencing (32,33,66,67). This phenomenon was also observed in GFP^+^-hDPSCs grafts, where one month after transplantation not all human-origin cells retained GFP expression. Some GFP-negative cells exhibited positive immunolabeling for the human-specific marker STEM121. Furthermore, GFP-expressing cells typically localized near the graft epicenter, consistent with observations from neural precursor grafts in adult spinal cord and striatum (67,68). These findings were corroborated by immunohistochemistry, showing that most GFP-retaining cells were DCX-positive, consistent with an immature neuroblast-like phenotype.

Development and functional integration of grafted cells within the host tissue generally require prolonged maturation periods. Embryonic stem cell-derived neural progenitors display immature electrophysiological properties at 6-12 weeks post-grafting and do not acquire mature firing patterns or spontaneous postsynaptic currents until 18-24 weeks (69–71). Similarly, induced pluripotent stem cell-derived neurons lack sustained firing capabilities until 12-50 weeks *in vivo* (40). Consistent with these findings, we observed an immature electrophysiological profile in GFP^+^-hDPSCs four weeks after hippocampal transplantation. In current-clamp mode, grafted hDPSCs generated single TTX-sensitive APs but failed to exhibit repetitive firing patterns. Comparable results have been reported for endogenous DCX-positive neural progenitors in the adult subventricular and subgranular zones (72).

Neuroblast’s electrophysiological characterization in acute brain slices results a technically complex process, which presents many limitations. Due to their small somatic size, patch clamp recordings are hindered by rapid dialyses of the intracellular content. This leads into a quick cell death in less than 10 minutes after making the seal. Furthermore, when brain slices are recorded close to physiological temperatures, neuroblasts migrate at a rate of approximately 50-60 μm/h with a consequent loss of the seal and the recordings (38). We could also experience these difficulties in our own recordings. Due to the small size of GFP^+^-hDPSCs only few recordings could be made after sealing the cell and before its death.

Immature neural cells exhibit functional properties that differ markedly from those of mature neurons. Generally, large and more mature neurons possess a high density of ion channels distributed over an expanded membrane surface, facilitating ionic current flow and resulting in lower input resistances, typically around 1GΩ. In contrast, neuroblasts express fewer voltage-gated ion channels and therefore display significantly higher input resistances, usually close to 4 GΩ (72,73). Consistent with these characteristics, the input resistance of grafted non-differentiated GFP^+^-hDPSCs measured on month after injection was approximately 2.4 GΩ.

In addition to developmental stage, pharmacological manipulation can importantly influence input resistance. Specifically, TTX application induces a reduction in input resistance by blocking voltage-gated sodium channels and consequently altering membrane conductance. In hippocampal pyramidal neurons, TTX treatment results in input resistance values approximately 22% lower than those observed under control conditions (74). A comparable reduction in input resistance was observed in grafted hDPSCs following TTX perfusion.

Beyond input resistance, immature neurons are further characterized by lower membrane capacitance values around 8 pF, whereas mature neurons typically show values close to 17 pF (75). In agreement with their yet immature neuronal phenotype, GFP^+^-hDPSCs grafted into the hippocampus displayed a mean capacitance of 13 pF.

Overall, our study provides the first evidence that neurally preconditioned hDPSCs derived from dentospheres can survive and integrate into the hippocampal network of non-immunosuppressed C57BL/6J mice. Nevertheless, the persistence of DCX-positive labeling together with immature, neuroblast-like electrophysiological properties observed at one-month post-transplantation, indicates a yet incomplete maturation. Therefore, future studies should include functional characterization at longer post-grafting time points.

## Acknowledgement

We would like to thank to Ricardo Andrade and Alex Díez from the Analytical and High Resolution Microscopy Service in Biomedicine of the SGIker services (EHU), Rafael Martínez Conde’s maxillofacial surgery clinic and Murielle Févre from the *Équipe du Pôle In Vivo Experimental* at the PIV-EXPE, Centre Broca Nouvelle-Aquitaine. This work has been funded by the University of the Basque Country (EHU) (grant EHU-G24/08) and EHUN24/54 to J.L, the Basque Government (IT1751-22; to G.I.; ELKARTEK program MYOZET KK-2024/00111 to G.I.; “Strengthening strategic health research” program No. 2023333035 to J.R.P.), grant PID2023-152704OB-I00 (J.R.P. and G.I.) funded by MCIN/AEI/10.13039/501100011033 and by the European Union (NextGenerationEU) “Plan de Recuperación Transformación y Resiliencia”, POLIMERBIO SL (EHU) contract 2023.0012). I.M.R. obtained a Ph.D. fellowship from University of the Basque Country (EHU) (PIFBUR21/05). B.P.R. obtained a post-doctoral fellowship from Basque Government (POS_2025_1_0061). J.S.M., D.M.A. and S.H.S. obtained a Ph.D. fellowship from Basque Government (Ref. PRE_2025_2_0087, PRE_2025_2_0049, PRE_2025_1_0157). The funding sources had no role in the study design, data collection, data analysis, data interpretation, writing of the manuscript, or decision to submit it for publication.

## Data availability statement

The data that support the findings of this study are available from the corresponding authors upon reasonable request.

## Author contributions

B.P.R., F.U., F.L., G.I. and J.R.P. were responsible for the study concept and design. B.P.R, I.M.R., J.S.M., R.B.T, D.M.A., S.H.S, J.L., M.A., and A.L.S.A., performed the investigation and formal analysis. B.P.R, G.I. and J.R.P. contributed to the methodology and writing of the original draft. F.L., F.U., G.I. and J.R.P. handled conceptualization, funding acquisition and supervision. All authors reviewed and critically revised the draft manuscript. Authors approved the final manuscript.

## Ethics approval

The study protocol was approved on date 03/02/2021 by the Ethics Committee of the University of the Basque Country (EHU) and the competent authority (Provincial Administration of Bizkaia) regarding the use of human cells with CEISH M10/2020/172 and M10/2023/025. The animal procedures were approved by the Ethics Committee of the University of the Basque Country (EHU) and the competent authority (Provincial Administration of Bizkaia) regarding the use of human cells and experimental animals with CEISH M10/2023/025 and CEEA M20/2023/026, M30/2023/027 and the ethics committees at the Interdisciplinary Institute of Neuroscience in Bordeaux and the competent authorities in France (project reference: 46334, dossier reference: 202312181625835).

## Declaration of competing interest

The authors report no biomedical financial interests or potential conflicts of interest.

## Abbreviations

aCSF: Artificial Cerebrospinal Fluid
AP: Anteroposterior
APs: Action Potentials
bFGF: Basic Fibroblast Growth Factor
BSA: Bovine serum albumin
CaCl_2_: Calcium chloride
CNS: Central nervous system
DMEM: Dulbecco’s Modified Eagle Medium
DV: Dorsoventral
EGF: Epidermal Growth Factor
ESCs: Embryonic stem cells
FBS: Fetal bovine serum
GFAP: Glial Fibrillary Acidic Protein
GFP: Green fluorescent protein
hDPSCs: Human dental pulp stem cells
HEPES: 4-(2-hydroxyethyl)-1-piperazineethanesulfonic acid
iPSCs: Induced pluripotent stem cells
KCl: Potassium chloride
L: Laterolateral
Mg-ATP: Magnesium Adenosine Triphosphate
MgCl_2_: Magnesium chloride
mOsm: Milliosmoles
ms: Milliseconds
mV: Millivolts
mΩ: Megaohms
NaCl: Sodium chloride
Na-GTP: Guanosine 5′-triphosphate sodium salt hydrate
NaH_2_PO_4_: Sodium dihydrogen phosphate
NaHCO_3_: Sodium bicarbonate
pA: Picoamperes
PBS: Phosphate buffered saline
PFA: Paraformaldehyde
PNS: Peripheral nervous system
rpm: Revolutions per minute
TTX: Tetrodotoxin
TU/ml: Transduction Units per ml

## Notes

### Competing Interest Statement

The authors have declared no competing interest.

